# Angle-resolved Measurements Reveal the Origin of Signal Anisotropy in Pump-probe Microscopy

**DOI:** 10.64898/2025.12.27.691690

**Authors:** Xiaomeng Jia, Warren S. Warren

## Abstract

Nonlinear optical microscopy modalities have become widely used in applications ranging from material characterization to tissue diagnosis. In complex systems, multiple mechanisms often contribute to the signal; for example, two-color pump-probe microscopy provides rich molecular signatures of the samples based on several nonlinear processes. Here we uncover a surprising complication, that in many cases the detection direction drastically alters the signal components and the overall amplitude. To understand the origin of this signal anisotropy, we first calculate and measure a simple case, the pump-probe signal of gold nanoshells at various detection angles, attributing the effect to scattering changes due to the sample refractive index change; we then extend this analysis, showing a striking directional dependence arising from distinct scattering profiles of two dominant nonlinear processes. These results demonstrate that comparing epi (backward) and transmission (forward) signals provides additional information, enabling cleaner separation of nonlinear contributions in pump-probe measurements.

---

Nonlinear optical microscopy has become an important tool in biological research and material science. Many nonlinear imaging modalities are based on one specific optical nonlinear process, such as multiphoton fluorescence ^1,2^, second harmonic generation (SHG) ^3^, stimulated Raman scattering ^4^, and coherent anti-Stokes Raman scattering (CARS) ^5^. A particularly useful nonlinear imaging technique developed in the past two decades is the pump-probe microscopy, a versatile, label-free, time-resolved modality that can characterize the malignant potential of early melanoma cases ^6^. In a pump-probe measurement, a pump laser pulse excites carrier populations in the sample and after a variable delay, a probe laser pulse is used to measure the change of absorption induced by the pump. The change in probe absorption strongly depends on carrier population transfer between states of the target molecules and thus provides molecular specificity and imaging contrast.

As a direct consequence of the absorption measurement, pump-probe can also access non-emissive nonlinear processes ^6^. For example, the two-photon absorption (TPA) and excited-state absorption (ESA), reduce the probe pulse when the pump pulse is on (our convention is to graph this as a positive signal). Other processes, such as ground state depletion (GSD) and stimulated emission (SE), strengthen the probe pulse when the pump pulse is on (by our convention a negative signal). Stimulated Raman scattering (SRS) can provide either sign, depending on the pump and probe wavelengths. The pump-probe signal thus is often a superposition of several nonlinear processes with various decay dynamics.

In nonlinear optical imaging, signals scale nonlinearly with incident intensity, leading to a localization of signal generation at the focus and allows for simple 3D imaging through scanning of the focal position ^7,8^. It is generally assumed that the signal character imprinted at the focus remains the same if the signal makes its way to the detector, either in transmission (forward) or epi (backward) detection. For example, multiphoton fluorescence signal has the same decay lifetime in both directions; CARS signal is expected to have the same Raman shift peaks in transmission- and epi-detection, despite that epi-CARS can suppress the solvent background ^9,10^. The scattering profiles of signal were well investigated for coherent processes like SHG and sum-frequency generation (SFG), which exhibited that these nonlinear signals are highly directional upon generation in spherical particles ^11–14^. In bio-tissue and colloid samples, nonlinear signals could be deflected to various directions after generation. In most cases, the common sense is to collect signals in directions with the highest signal to noise ratio (SNR).

It came as a surprise to us, then, when we observed that pump-probe readout of some samples often appears qualitatively different in the transmission and epi detections. We realized that nonlinear signal scattering in incoherent nonlinear processes involving transfer of carrier populations (those characterized by both signal intensity and decay lifetime) was barely reported. This motivated us to carefully study the angle-resolved signal in pump-probe measurements, using the setup shown in Fig. 1 (Methods).

**Figure 1.**
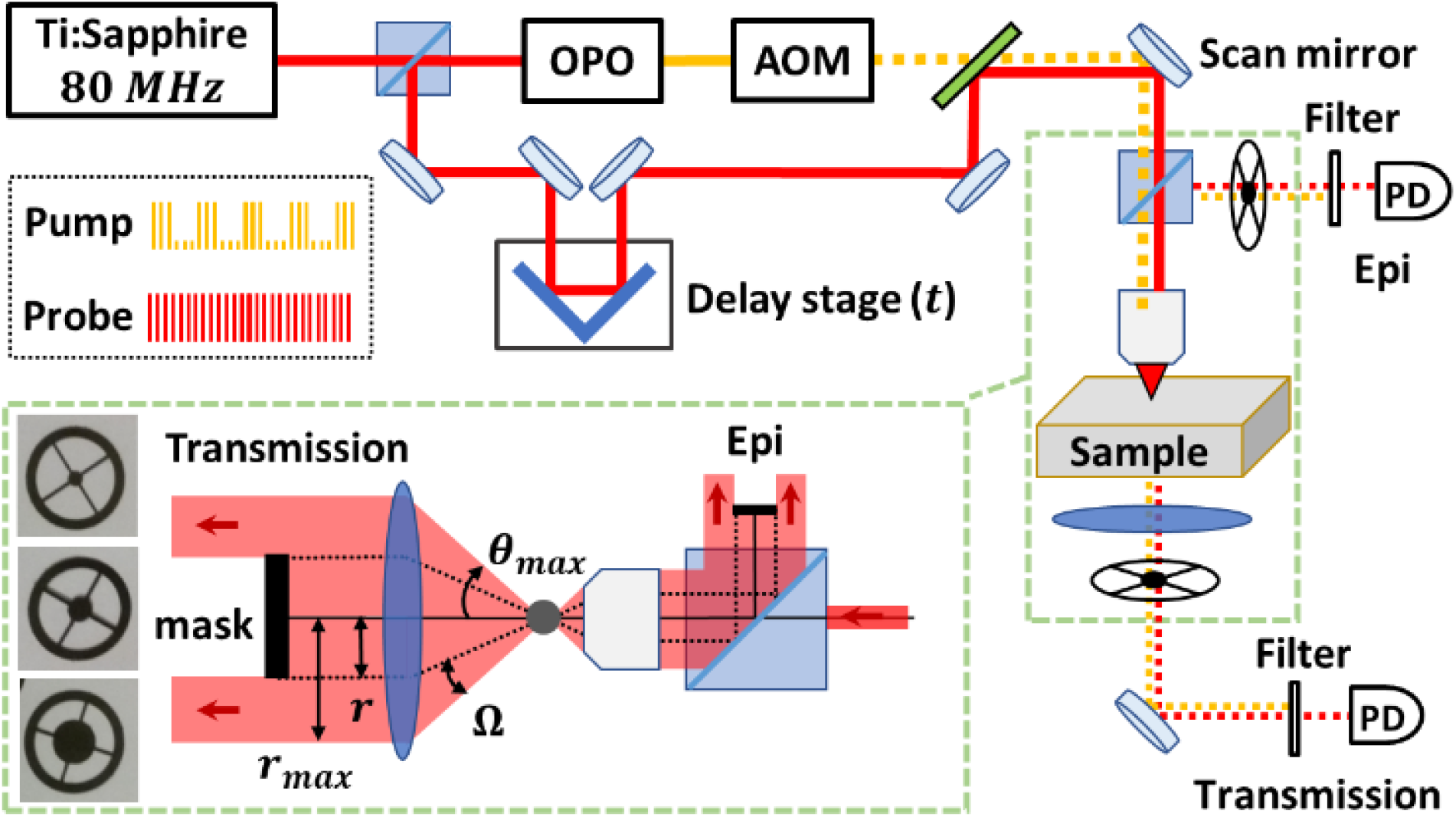
Setup of the pump-probe microscope with angle-resolved transmission and epi detections Pump (yellow) and probe (red) laser pulses are generated and overlapped at the sample focus to generate the pump-probe signal (Methods). To achieve angle-resolved detections, a mask of various sizes is mounted in either the transmission or epi detection pathway to block the central part of the signal, so that the pump-probe signal can be collected within a tunable solid angle Ω = *θ*_*min*_ − *θ*_*max*_. The upper limit *θ*_*max*_ is given by the NA of the optical condenser in the transmission detection (or by the NA of the objective in the epi detection), and the lower limit *θ*_*min*_ is tuned by the mask size.

## Results

Pump-probe microscopy shows the promise to grade metastatic potential of melanoma cases by detecting the nonlinear signatures of melanin pigment ^15–17^. But a striking feature is that thin melanoma biopsies (as well as natural and synthetic melanin particles) imaged in transmission detection usually produce signals consisting of both a negative GSD and a positive ESA (and a minor SRS at some wavelengths) ^18,19^. However, we notice that imaging in epi detection systematically exhibits only the ESA component. Fig. 2(a) shows typical pump-probe delay traces measured in mouse melanoma samples (Methods) in the two detection directions. This transmission-epi discrepancy is not due to heterogeneity of the melanin particles, as measurements on synthetic melanin particles of a uniform size (Methods) show similar observations (Fig. S1(a) in Supplementary). The discrepancy is also observed in many other materials, such as the synthetic ultramarine – the main content of the blue pigment lapis lazuli used in historical artworks (Fig. S1(b) in Supplementary).

**Figure 2.**
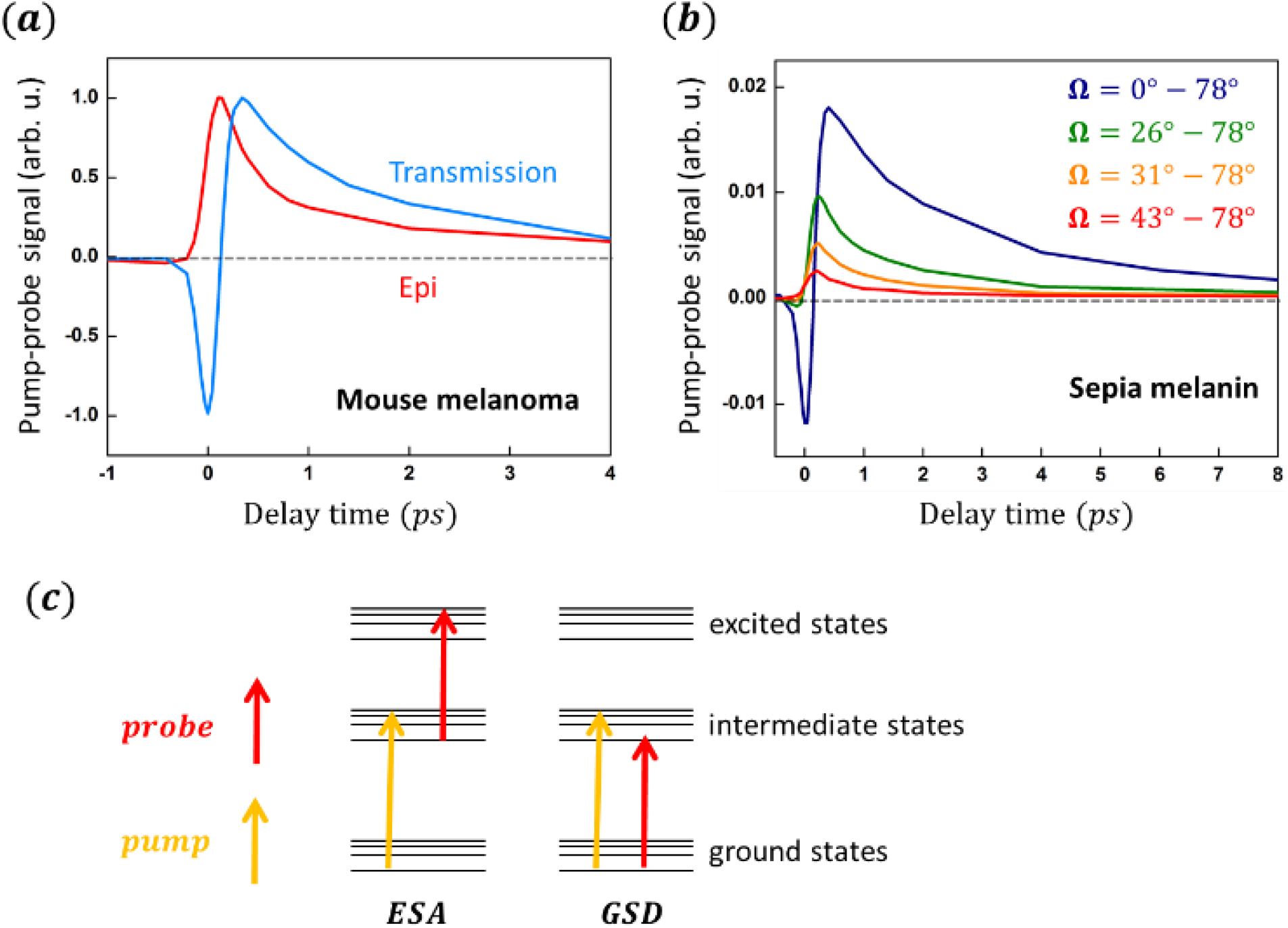
Transmission-epi discrepancy of pump-probe signal in melanin samples and illustration of the underlying mechanism from two contributing non-linear processes (***a***). Mouse melanoma samples imaged in transmission and epi detections have different delay traces. Pump/probe wavelength: 720 *nm*/817 *nm*. Both curves are normalized at peak positive amplitudes for comparison. (***b***). Angle-resolved measurements in transmission detection on sepia melanin particles indicate that the negative GSD signal is observable only in the forward direction. Pump/probe wavelength: 743 *nm*/817 *nm*. (***c***). Electronic energy diagrams of ESA and GSD. The probe beam detects the electron population of excited states in ESA and the hole population of ground states in GSD.

To narrow down the angular range where the signal changes, we image natural melanin particles from sepia officinalis (Methods) with the setup shown in Fig. 1, where signals within a tunable solid angle Ω = *θ*_*min*_∼*θ*_*max*_ can be collected in either detection. As Fig. 2(b) shows, the GSD vanishes in off-axis angles, while ESA is observable at various angle ranges. Thus, it seems that GSD is carried only in the incidental direction. As sepia melanin contains various particle sizes with consequently different scattering profiles, we verify the generality of these features using synthetic melanin particles of defined sizes (Fig. S2, S5, S6, Supplementary) and at various pump/probe wavelengths (Fig. S3, Supplementary), showing the same directional dependence of ESA and GSD. Angle-resolved measurements in epi detection on synthetic melanin particles confirm that the ESA is always observable, while GSD never appears, in various back-scattering angles (Fig. S4, Supplementary). Thus, the directional nature of GSD underlies the transmission-epi discrepancy in melanoma imaging. To illustrate the different directional dependence of the GSD and ESA, their electronic energy diagrams are shown in Fig. 2(c). Although neither of them is a coherent nonlinear process, GSD requires complete depletion of the ground-state electrons, necessitating sufficiently high pump power along the detecting axis. In contrast, ESA depends on excited-state electron populations that can be generated by the pump regardless of its incident direction. Thus, the probe transitions in ESA and GSD correspond to distinct scattering and absorption coefficients, leading to different angular profiles of the collected signals.

Based on our analysis above, it is possible to calculate the detailed angular profile of pump-probe signals if the change in sample absorption and scattering coefficients due to pump excitation can be analytically derived. Given the featureless linear absorption spectrum and complex pump-probe signal compositions (GSD and ESA) in melanin particles, we decide to demonstrate this calculation of the pump–probe signal’s angular distribution in a simple system: gold nanoshell particles. The principle is to compute the change in the dielectric function Δ*ϵ* of gold upon pump excitation, use it to obtain the pump-probe spectrum, and then determine the angular distribution of signal at a given probe wavelength.

In gold, the pump-probe delay traces reveal the transient process of electron gas heating and cooling ^20^. When gold is excited by an inferred pump pulse, the electron gas in 6*s* band goes through a thermalization process to reach a quasi-equilibrium (Fermi-Dirac distribution at a higher temperature). The change in electron gas temperature leads to a change in *ϵ*. The calculation of Δ*ϵ* was well established in literature ^21,22^. Pump-probe spectrum of a gold nanoparticle at peak intensity is calculated by:

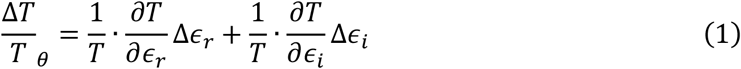

where *θ* is the detection angle with respect to the incident direction; *ϵ*_*r*_ and *ϵ*_*i*_ are the real and imaginary part of the dielectric function in gold; Δ*ϵ*_*r*_ and Δ*ϵ*_*i*_ are the real and imaginary part of Δ*ϵ* caused by pump laser; the coefficients 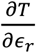 and 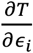 are dependent on the geometry of the nanoshell particles. The approach in literature ^23,24^ included only the absorption cross section in calculating 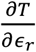 and 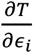, as scattering is negligible for small particles. In our calculations, we include both absorption and scattering cross sections.

To demonstrate the signal anisotropy origin, we are especially interested in studying the pump-probe signal carried on scattered probe light, so the ideal model sample should have a strong scattering and weak absorption (i.e., *σ*_*abs*_ ≪ *σ*_*sca*_) in the resonance of the pump-probe spectrum. Based on the criterion, a *SiO*_2_/*Au* nanoshell sample of particle size 242 *nm* and shell thickness 22 *nm* is selected (Methods). The sample has a size distribution of 242 ± 12 *nm* (Fig. S7, Supplementary). As shown in the calculated cross sections and pump-probe spectra at the incident angle (Fig. 3), the signal resonances are near 700 *nm*. At 720 *nm*, the scattering cross section *σ*_*sca*_ is about ten times larger than the absorption cross section *σ*_*abs*_, which makes it an ideal choice.

**Figure 3.**
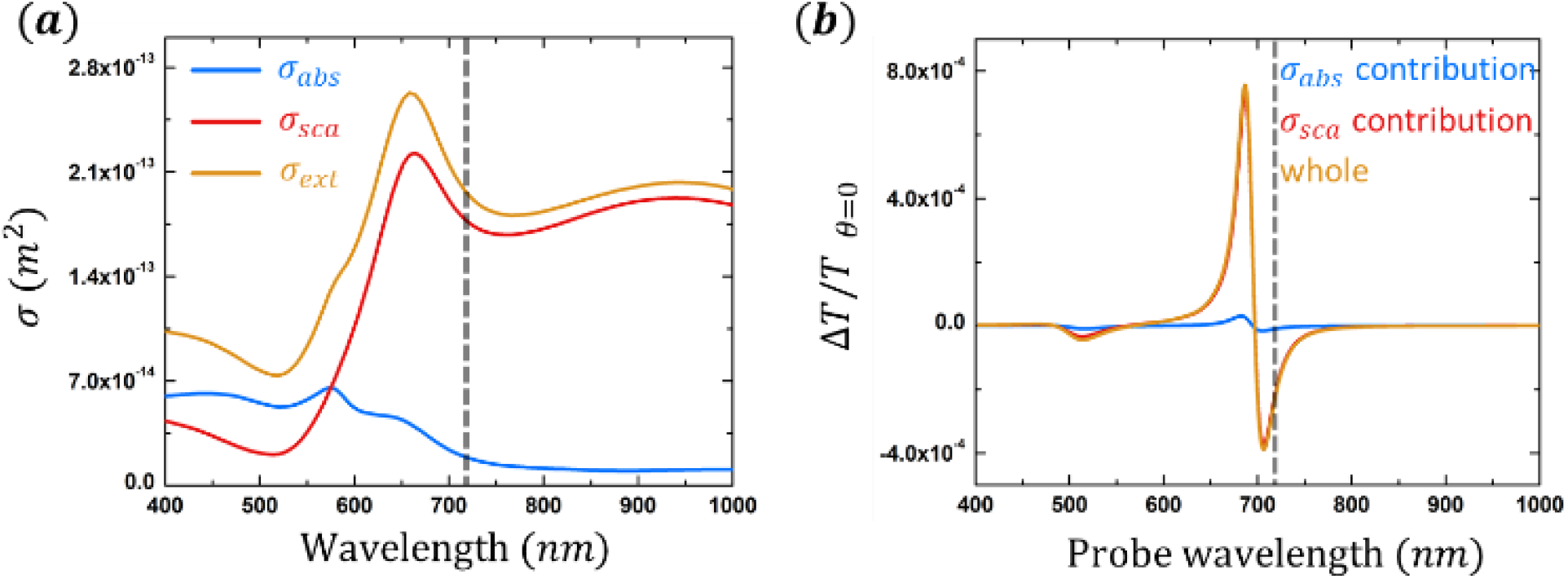
Calculated absorption/scattering cross sections and spectra of the gold nanoshell sample (***a***). Calculation shows that the scattering cross section *σ*_*sca*_ dominates at 720 *nm* (dash line) for the 242 *nm Au*/*SiO*_2_ nanoshell particle with shell thickness 22 *nm*. (***b***). Calculated pump-probe spectra of the gold nanoshell sample at detection angle *θ* = 0°. At probe wavelength 720 *nm* (dash line), contribution from scattering is about ten times stronger than from absorption, making it an ideal candidate to study the scattering-related pump-probe signal.

The angular distribution of pump-probe signal Δ*T*_*θ*_ at pump 817 *nm* and probe 720 *nm* was calculated and measured for the nanoshell sample (Fig. 4a). We used Mie scattering theory ^25–27^ to calculate the scatterings of probe laser at pump off and pump on. The difference between the two scattering profiles gives the angular distribution of the pump-probe signal. The calculation parameters are given in Supplementary.

**Figure 4.**
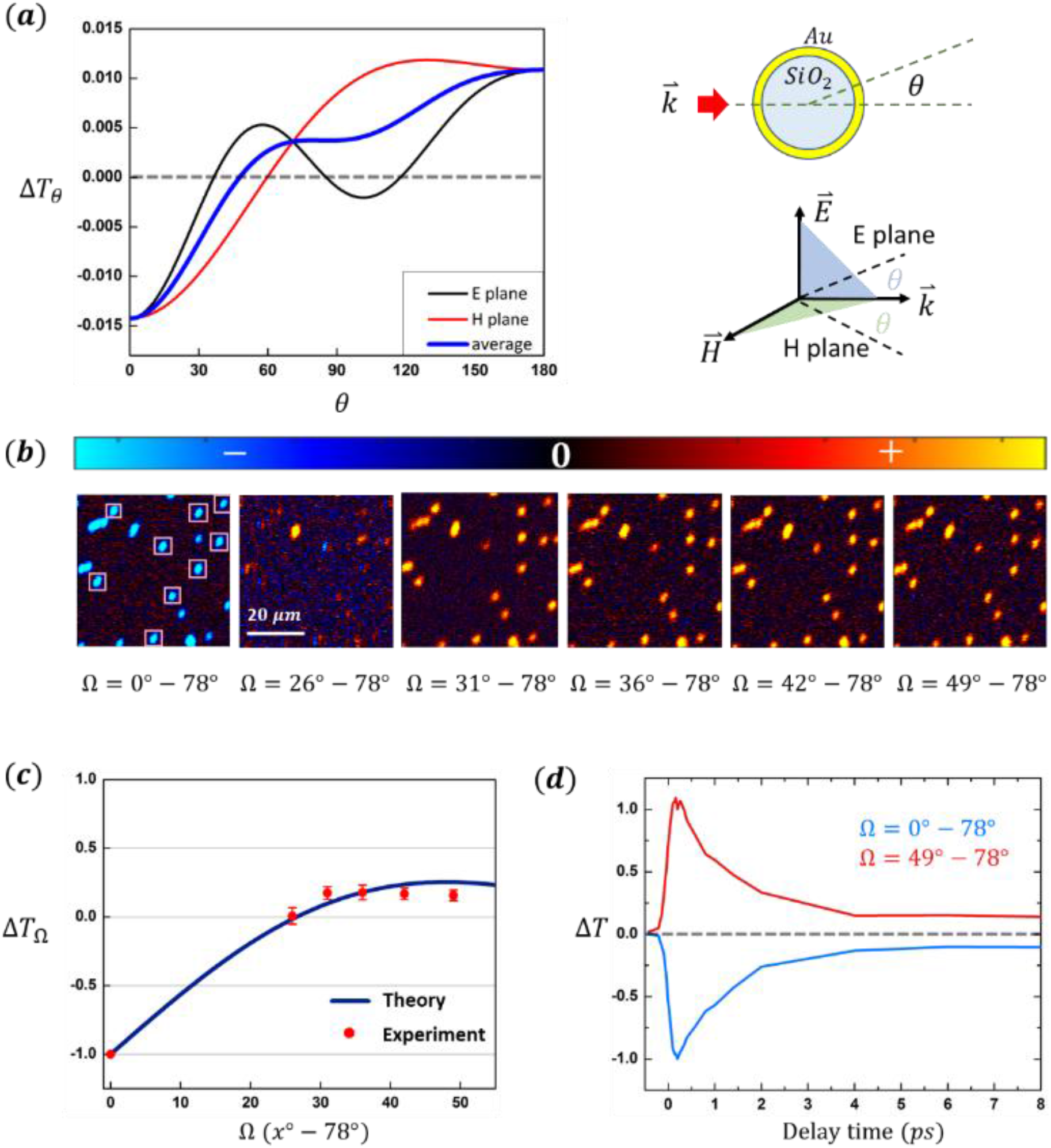
Pump-probe signal anisotropy demonstrated using the *Au*/*SiO*_2_ nanoshell particles (***a***). Calculated angular profiles of the pump-probe signal at *t* = 0.2 *ps* with probe wavelength 720 *nm*. The blue curve is the average of the *E* plane and *H* plane signals. The transition from negative to positive sits at *θ* = 49°. (***b***). At different detection solid angles Ω, measured pump-probe images of the 242 *nm* gold nanoshells with pump/probe = 817 *nm*/720 *nm* and delay time *t* = 0.2 *ps*. The incident laser power is 1.78 *mW*/2.31 *mW*. The images are false colored to indicate the signal sign (blue indicates negative and red/yellow indicates positive signal). (***c***). Measured signal strength at *t* = 0.2 *ps* for the 9 individual particles marked in (b) compared with calculation results (integration of the blue curve in (a)). Error bars are SD. The transition from negative to positive signal is clear. (***d***). The normalized delay curves at detection angles Ω = 0°∼78° and Ω = 49°∼78° show the identical decay dynamics (both normalized at intensity at *t* = 0.2 *ps*).

As the calculated Δ*T*_*θ*_ shows (Fig. 4(a)), although the signal distributions are different in E plane (formed by 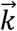 and 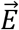 of incident probe beam) and H plane (formed by 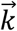 and 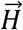 of incident probe beam), they both exhibit a sign-flip from negative to positive when detection angle *θ* increases. The measurable quantity Δ*T*_Ω_ is an integration of the Δ*T*_*θ*_ from *θ*_*min*_ to *θ*_*max*_ = 78°. Fig. 4(b) shows the images of the gold nanoshells at delay time *t* = 0.2 *ps* at several solid angle Ω values. The transition from negative to positive signal as a function of Ω is displayed in the images. The measured Δ*T*_Ω_ is consistent with calculation results (Fig. 4(c)). The decay dynamics measured at Ω = 0°∼78° and Ω = 49°∼78° are identical except for the sign-flip (Fig. 4(d)).

## Discussion

In this work, we made a surprising discovery that the pump-probe signal collected in the transmission and epi detections are qualitatively different in melanoma imaging (Fig. 2). For the first time, we reported the development of an angle-resolved detection method to investigate this phenomenon (Fig. 1). We found that the two signal components in melanin measurements, ESA and GSD, have different angular profiles (Fig. 2). We propose that this difference could be due to the distinct absorption and scattering coefficients related to relevant electron populations in the two nonlinear processes. As a demonstration, we validated the assumption regarding absorption and scattering coefficients underlying signal anisotropy using a simple, calculable gold nanoshell sample. The calculations and measurements on the gold nanoshell samples showed how the dielectric function *ϵ*, which contains information of absorption and scattering coefficients of the particle, leads to the angular anisotropy in pump-probe signal via scattering (Fig. 3, 4).

Significantly, this discovery indicates that detection angles could serve as another parameter for pump-probe imaging. Some nonlinear processes (like GSD in this case) are highly dependent on detection angles, regardless of particle size and over a wide range of laser wavelengths, while others are not. This work presents the first implementation of an angle-resolved detection method in pump-probe microscopy, based on which we can decompose the signal and fit their decay dynamics separately, potentially improving the data analysis of pump-probe imaging. This state-of-the-art method can be adopted by other laboratories for many future research topics.

## Acknowledgment

We thank Dr. S. Degan for providing the mouse melanoma samples. We thank Dr. D. Grass and Dr. M. Fisher for helpful discussions and comments on the manuscript. This work is supported by the National Institutes of Health grant R01CA166555 and National Science Foundation Grant CHE-1610975 (to W. S. W.). Some figures in this manuscript are adapted from the author’s doctoral dissertation (Jia, 2021) ^28^.

## Author Contributions

X. J. designed the assays, prepared all the samples, updated the pump-probe instrument, and performed all the calculation, measurements, and data analysis. W. S. W. supervised the project. X. J. and W. S. W. wrote the initial draft.

## Data Availability Statement

All data supporting the findings in this paper are provided in the main manuscript and its Supplementary file.

