## Supplementary Information for "Angle-resolved Measurements Reveal the Origin of Signal Anisotropy in Pump-probe Microscopy"

### Methods

#### Samples

##### Preparation of mouse melanoma samples

A B16 melanoma mouse model was used for this study. The mouse was born on April 18, 2017. On June 16, 2017, B16 melanoma cells were injected subcutaneously. The animal was identified by a distinctive large hole in the right ear. Treatment with Trametinib (0.6 mg/kg) was initiated on June 29, 2017 and administered daily. On July 3, 2017, the melanoma tumor measured 5.5 × 6.5 mm. The tumor appeared black on the surface and exhibited no edema. Following euthanasia, the tumor was dissected, placed in phosphate-buffered saline (PBS), and mounted on a glass slide for imaging.

##### Preparation of Sepia melanin particles

Melanin from Sepia officinalis was purchased from Sigma-Aldrich (CAS Number: 8049-97-6) as a powder, ground with a mortar into a finer powder, and dispersed in deionized (DI) water. The melanin colloid was purified by repeated centrifugation ($1000\ rpm$) and redispersion in DI water until all visible aggregates were removed. A 2% agarose solution was prepared by mixing agarose powder (0.2 g) with DI water (10 g) and heating until the mixture became transparent. The purified melanin colloid was then combined with the agarose solution at equal volumes, and the mixture was deposited into a $\sim 70\ \mu m$ thick groove on a glass slide and covered with a coverslip. Upon cooling to room temperature, the agarose solution solidified into a gel, immobilizing the sepia melanin particles within the matrix. Freshly prepared samples were imaged using the pump-probe microscope.

##### Preparation of synthetic melanin particles

DHI-type (5,6-dihydroxyindole) melanin particles were prepared following a previously reported protocol [1]. To synthesize 300 nm melanin particles, dopamine hydrochloride (180 mg; Sigma-Aldrich, CAS No. 62-31-7) was dissolved in $90\ mL$ DI water. At room temperature and under vigorous stirring, $NaOH$ solution ($650\ \mu L$, $1N$) was added to the dopamine hydrochloride solution. The mixture was stirred continuously for $6\ h$, during which the solution gradually turned dark brown. The synthesized melanin particles were collected by centrifugation ($5000\ rpm, 15\ min$) and washed several times with DI water until the supernatant became clear.

To synthesize $100\ nm$ melanin particles, dopamine hydrochloride ($180\ mg$) was dissolved in $90\ mL$ of DI water. Under vigorous stirring at $50$℃, $NaOH$ solution ($770\ \mu L, 1N$) was added to the dopamine hydrochloride solution. The mixture was maintained at $50$℃ with continuous stirring for $6\ h$, during which the solution gradually turned dark brown. The synthesized melanin particles were collected by centrifugation ($20000\ rpm, 15\ min$) and washed with DI water several times until the supernatant became clear.

Sizes of the synthesized melanin particles were confirmed using scanning electron microscopy (SEM) and transmission electron microscopy (TEM) (see Fig. S5 and S6).

The synthesized melanin solutions were mixed with $2\%$ agarose solution at equal volume. The mixture was deposited into a $\sim 70\ \mu m$ thick groove on a glass slide and covered with a coverslip. The agarose solution solidified into a gel upon cooling down to room temperature, immobilizing the melanin particles within the matrix. The freshly prepared samples were then imaged using the pump-probe microscope.

**Preparation of gold nanoshell particles**

Gold nanoshell particles ($Au/SiO_2$) with a total diameter of $242\ nm$ and a shell thickness of $22\ nm$ were purchased from nanoComposix (GSPN980-25M) and diluted in DI water. The particle suspension was mixed with $2\%$ agarose solution at equal volumes, deposited onto a glass slide, and covered with a coverslip. The agarose solution solidified into a gel to immobilize the nanoshell particles within the matrix. The freshly prepared samples were then imaged using the pump-probe microscope.

#### Apparatus

**Pump-probe microscopy**

As shown in Fig. 1, a mode-locked Ti:Sapphire laser (Coherent, Chameleon) generated ultrashort pulse trains with a tunable wavelength and a repetition rate $80\ MHz$. The laser beam was split into two paths. One beam was directed through an optical parametrical oscillator (OPO; Coherent, Mira-OPO) to change its wavelength. The intensity of the OPO beam was modulated at $2\ MHz$ using an acousto-optic modulator (AOM) and served as the pump beam, while the other beam served as the probe. The delay time ($t$) between the two beams was controlled by adjusting the relative optical length using a motorized translational stage (delay stage).

**Scanning Electron Microscopy (SEM)**

Synthetic melanin particles were characterized using a field-emission scanning electron microscope (FEI XL30 SEM-FEG) operated at an accelerating voltage of 30 kV with a working distance of 6.8 mm.

**Transmission Electron Microscopy (TEM)**

Synthetic melanin particles were characterized using a transmission electron microscope (FEI Tecnai $G^2$ Twin).

#### Material parameters in scattering calculation

The parameters used in the calculation of the nanoshell sample include: refractive index of medium (water) at $720\ nm$, $n_{medium} = 1.33$; refractive index of core (silica) at $720\ nm$, $n_{core} = 1.43$; refractive index of shell (gold) at $720\ nm$ before pump excitation, $n_{shell}(off) = 0.169603101045296 + 4.05622799070848i$ and after pump excitation, $n_{shell}(on) = 0.17277678272209 + 4.04985051918599i$; core radius $r_1 = 99\ nm$ and particle radius $r_2 = 121\ nm$, and probe wavelength $\lambda = 720\ nm$.

The $n_{shell}(off)$ value was given by the Brendel-Bormann model [2], and corresponds to the dielectric function of gold at $720\ nm$, $\epsilon_{off}(720\ nm) = -16.4242203007228 +$

$1.37589769154178i$, $n_{shell}(on)$ was converted from $\epsilon_{on}(720\ nm) = \epsilon_{off}(720\ nm) + \Delta\epsilon(720\ nm) = -16.3714374111033 + 1.39944028642069i$.

#### Supplementary Figures

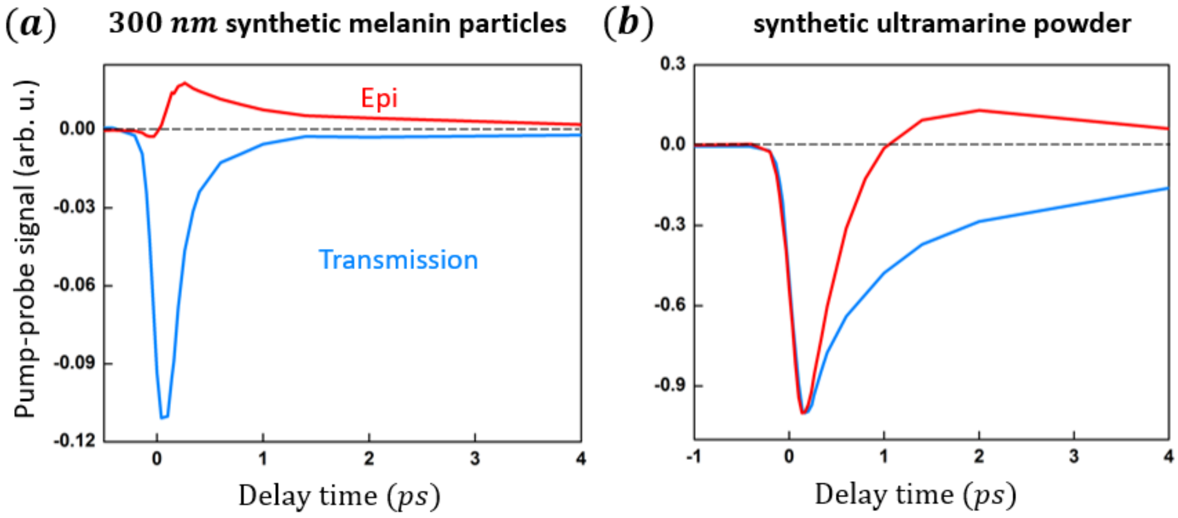


**Fig. S1** Pump-probe delay traces of (a) 300 $nm$ synthetic melanin samples and (b) synthetic ultramarine powders measured at pump 720 $nm$ and probe 817 $nm$ in transmission and epi detection mode. For both samples, the dominant nonlinear processes are GSD and ESA. Incident power was 1 $mW/1\ mW$ for $\boldsymbol{a}$ and 1.5 $mW/1.5\ mW$ for $\boldsymbol{b}$, respectively. The curves were averaged over $256 * 256$ pixels (imaging area of $144\ \mu m * 144\ \mu m$) in (a), and $128 * 128$ pixels ($360\ \mu m * 360\ \mu m$) in (b).

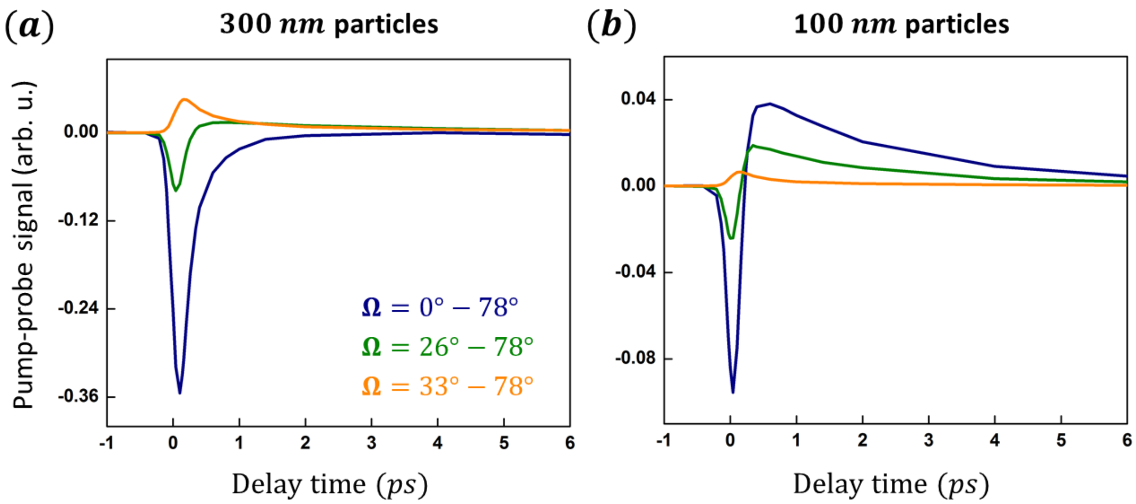


**Fig. S2** Angle-resolved measurements in transmission detection on synthetic melanin samples. Pump-probe delay traces of (***a***) 300 $nm$ and (***b***) 100 $nm$ synthetic melanin particles measured at pump 720 $nm$ and probe 817 $nm$ at different detection angles $\Omega$. Incident power was 1 $mW/1\ mW$. All the curves were averaged over $256 * 256$ pixels (imaging area of $144\ \mu m * 144\ \mu m$).

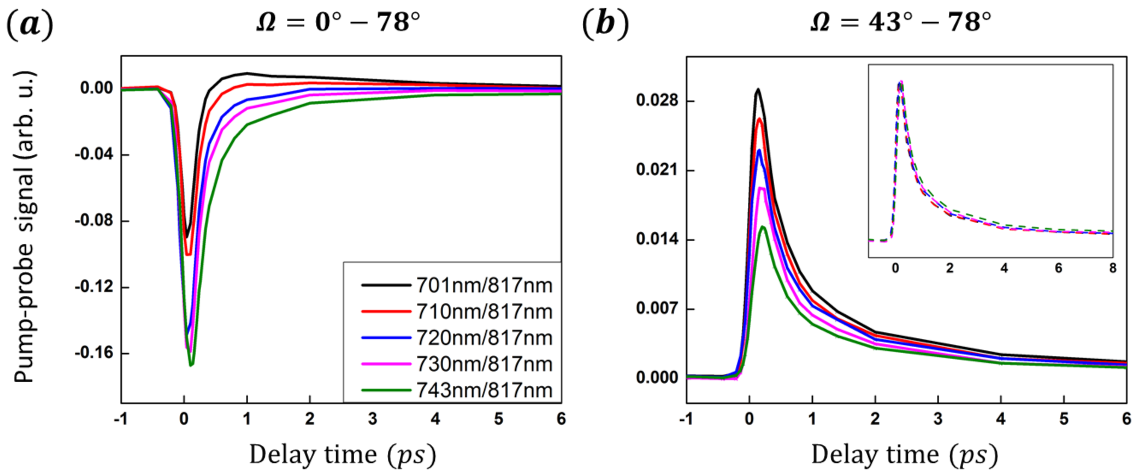


**Fig. S3** Pump wavelength sweeping measurements in transmission on synthetic melanin samples. Pump-probe delay curves of 300 $nm$ synthetic melanin particles measured at pump wavelength $701, 710, 720, 730, 743$ $nm$ and probe wavelength 817 $nm$ at detection angles (***a***) $\Omega = 0° - 78°$ and (***b***) $\Omega = 43° - 78°$. The inset in (***b***) is normalized delay traces. Incident power was $0.6\ mW/0.6\ mW$. All the curves were averaged over $256 * 256$ pixels (imaging area of $36\ \mu m * 36\ \mu m$).

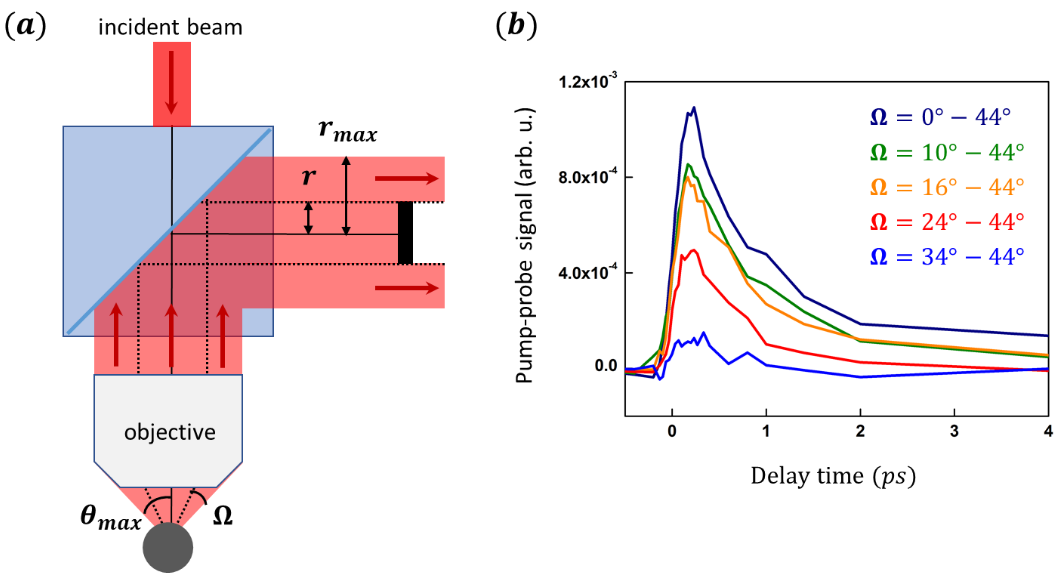


**Fig. S4** Angle-resolved measurements in epi detection on synthetic melanin. (***a***) Diagram of epi detection within a tunable solid angle $\Omega = \theta_{min} - \theta_{max}$. The upper limit $\theta_{max} = 44°$ was restricted by the objective (Olympus UPlanApo, $NA = 0.7$). The lower limit $\theta_{min}$ was tuned by the mask size. The divergency of the incident probe beam (817 $nm$) was ~8° as calculated by the focal length (9 $mm$) and beam size (~2.5 $mm$) at the objective pupil. (***b***) The delay traces of 300 $nm$ synthetic melanin particles measured at pump 700 $nm$ and probe 817 $nm$ at various epi detection angles $\Omega$ only contain the positive ESA. Incident power was $1\ mW/1\ mW$. All the traces were averaged over 8 measurements over $256 * 256$ pixels (imaging area of $72\ \mu m * 72\ \mu m$).

(*a*) (*b*)

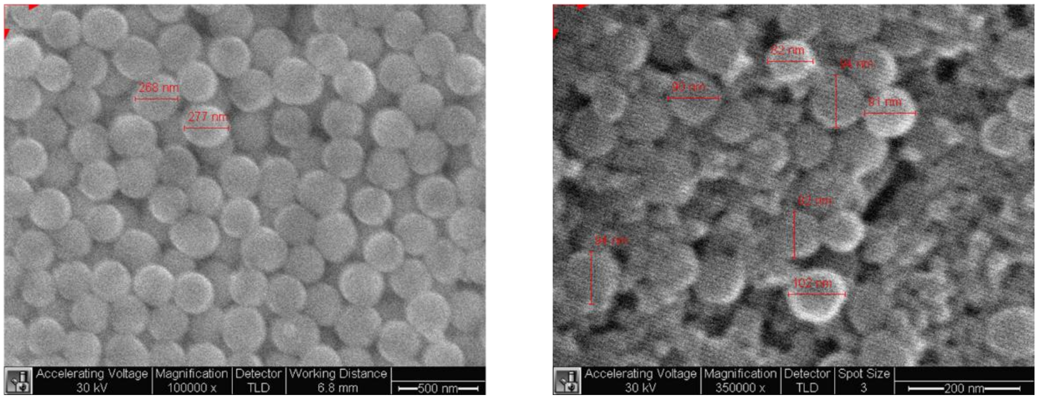


**Figure S5** SEM images of (a) $300\ nm$ and (b) $100\ nm$ synthetic melanin particles.

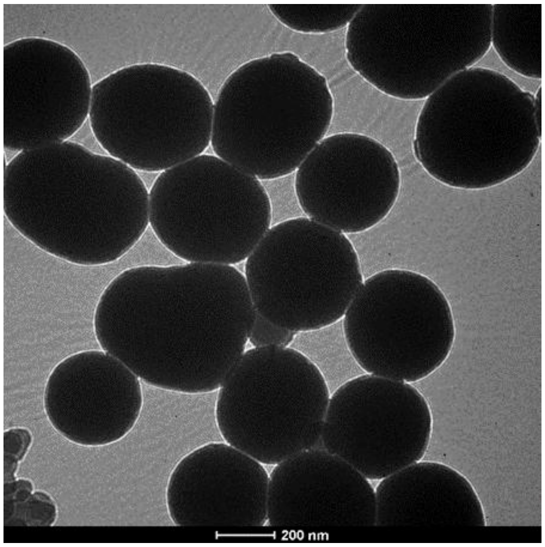


**Figure S6** TEM image of $300\ nm$ synthetic melanin particles.

Lot Number: ALJ0120

| | | | |
|---|---|---|---|
| **Total Diameter ± Std.Dev (TEM):** | 242 ± 12 nm | **Mass Concentration (Au):** | 0.053 mg/mL |
| **Coefficient of Variation:** | 4.8 % | **Hydrodynamic Diameter:** | 283 nm |
| **Core Diameter (TEM):** | 198 ± 10 nm | **Zeta Potential:** | -40 mV |
| **Shell Thickness (Calc'd):** | 22 nm | **pH of Solution:** | 5.6 |
| **Surface Area (Calc'd):** | 2.5 $m^2/g$ | **Particle Surface:** | PVP 40 kDa (Polymer) |
| **Particle Concentration (Calc'd):** | 8.1E+08 particles/mL | **Solvent:** | Milli-Q Water |

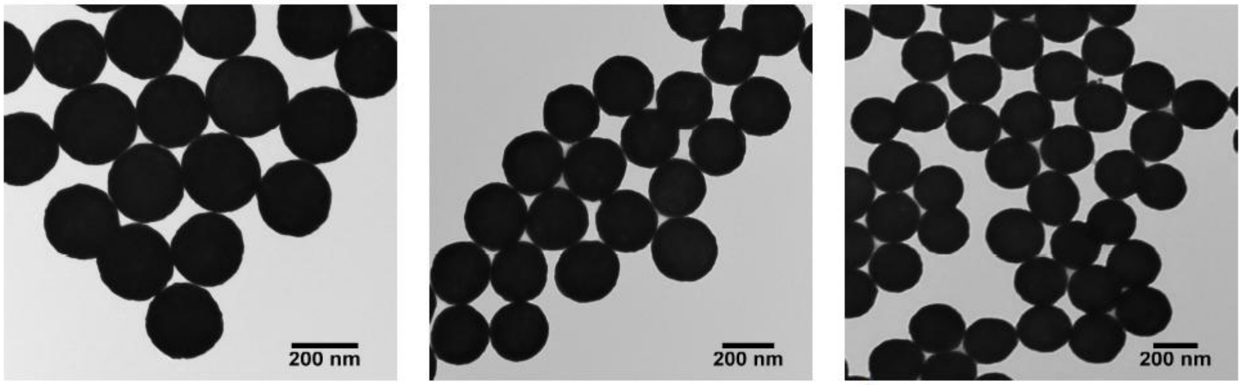


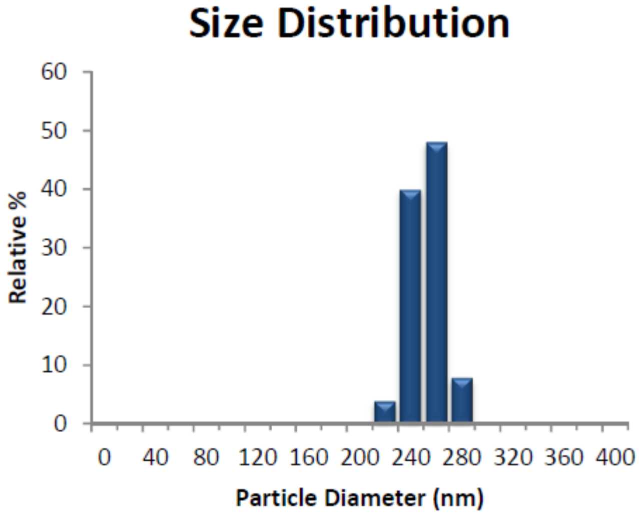


**Fig. S7** Certificate of Analysis for the $242\ nm$ gold nanoshell sample.
